# A TCR-mimic bispecific antibody reduces HIV-1 provirus and delays viral rebound in HLA-matched humanized mice

**DOI:** 10.1101/2025.10.06.680344

**Authors:** Zhe Yuan, Nathan L. Board, Miaoyun Zhao, Guorui Zu, Srona Sengupta, Qingsheng Li, Janet D. Siliciano, Robert F. Siliciano, Luis J. Montaner

**Affiliations:** The Wistar Institute, Philadelphia, PA, USA; Department of Medicine, The Johns Hopkins University School of Medicine, Baltimore, MD, USA; Nebraska Center for Virology, School of Biological Sciences, University of Nebraska-Lincoln, Lincoln, NE, USA; Howard Hughes Medical Institute, Baltimore, MD, USA

## Abstract

Bispecific antibodies that reroute cytotoxic effectors toward infected cells are promising HIV-1 cure agents, yet existing formats bind Env and are limited by antigenic variation and Env down-regulation. We engineered a TCR-mimic single-chain diabody, HI12, that recognizes a conserved Pol-derived peptide presented by HLA-A^*^02:01 and evaluated its effect in HLA-matched, HIV-infected humanized mice. When administered during early antiretroviral therapy (ART), HI12 was well tolerated, activated HIV-specific CD8^+^ T cells and accelerated plasma virus decay. Treatment produced four-to six-fold reductions in intact and total proviral DNA within lymph-node and splenic CD4^+^ T cells, indicating substantive reservoir clearance. After ART interruption, HI12-treated animals showed a significant delay in viral rebound compared with controls, linking reservoir reduction to improved post-therapy control. These findings provide the first in-vivo evidence that a peptide-HLA-directed bispecific antibody can both shrink the intact HIV reservoir and defer viral recrudescence, supporting further development of TCR-mimic bispecific antibodies for cure strategies.

**Significance Statement:** An HIV-1 cure strategy will require novel therapeutics to facilitate immune-mediated elimination of infected cells and reduction of blood and tissue reservoirs. Here, we demonstrate that an TCR-mimic bispecific antibody that recognizes a conserved Pol epitope can promote CD8^+^ T cell-mediated clearance of infected cells in HLA-matched, HIV-infected humanized mice. The bispecific antibody therapy reduced HIV-1 proviral DNA in lymph node and splenic tissues and improved post-therapy viral control. This study highlights the potential of developing novel TCR-mimic bispecific antibodies in HIV cure-directed strategies.

## Introduction

Despite effective suppression of viral replication through ART^1,2^, latent HIV-1 proviruses persist in long-lived memory CD4^+^ T cells^3,4^, necessitating lifelong treatment adherence. Although this population of HIV-1 reservoir cells decays slowly over the first seven years of ART^5^, recent studies indicate that reservoir decay does not continue and gradually increases after decades of treatment due to proliferation of latently infected cells^6,7^. Sustained viral control in the absence of ART may require substantial reduction of reservoir size in combination with boosted HIV-1-specific immune activity. Therefore, therapeutic interventions that facilitate reservoir cell elimination are greatly needed.

Bispecific antibodies have been proposed as one potential therapeutic strategy for facilitating HIV-1 reservoir reduction. By crosslinking CD8^+^ T cells or NK cells to HIV-1-infected target cells, bispecific antibodies are capable of promoting potent HIV-1-specific cytolysis^8–13^. We previously described an approach involving isolating single-chain variable fragments (scFvs) that recognize HIV-1 peptide:MHC complexes with high specificity and using the resulting scFvs to generate bispecific T cell-engaging antibodies^12^. This approach has also been employed in the development of TCR-mimic bispecific antibodies that recognize p53 and RAS neoepitopes for cancer immunotherapy^14,15^. In our prior study, we found that TCR-mimic bispecific antibodies possess potent HIV-1-specific CD8^+^ T cell activation and viral suppression activity in vitro^12^. Here, we evaluate the impact of early intervention with one of our previously described HIV-1-specific TCR-mimic bispecific antibodies when added during early ART in a bone marrow-liver-thymus (BLT) humanized mouse model of HIV-1 infection. We show that treatment with this bispecific antibody results in elevated CD8^+^ T cell activation and faster time to viral suppression in vivo. Furthermore, we find that the TCR-mimic bispecific antibody also promotes enhanced clearance of HIV-1-infected cells as evidenced by reduced HIV-1 DNA tissue burden and a delayed time to viral rebound upon ART interruption.

## Results

### In vivo administration of an HIV-1-specific TCR-mimic scDb is tolerable and induces CD8^+^ T cell activation

We independently generated two cohorts of humanized mice, each reconstituted from a different HLA-A^*^02:01^+^ tissue donor (n=22 for cohort I, n=21 for cohort II) (Figure 1A,B). Two weeks after infection with a transmitted/founder (T/F) HIV-1 strain (HIV_SUMA_), the mice were placed on antiretroviral therapy (ART). The mice were then randomly assigned to receive treatment with one of two TCR-mimic single-chain diabodies (scDbs) specific for different peptides in complex with HLA^*^02:01. H2 scDb (n=10 for cohort I, n=10 for cohort II) recognizes a human cancer p53 neoantigen-derived peptide and serves as a non-HIV-1-specific control^12,14^. HI12 scDb (n=12 for cohort I, n=11 for cohort II) recognizes a highly conserved HIV-1 reverse transcriptase (RT)-derived peptide that has previously been described as a target of HLA-A^*^02:01-restricted CD8^+^ T cell responses in PWH^12^. Both scDbs induce CD3-mediated T cell activation and cytolytic activity in the presence of cells presenting their target peptide in complex with HLA-A^*^02:01^12,14^. After one week of ART, the mice were administered the scDbs as daily 200μg intraperitoneal (IP) injections for two weeks (i.e., from day 21-35 postinfection or day 7-28 after ART start, Figure 1A,B). scDb IP injections were chosen to be administered daily due to the short plasma half-life (3.9 h) of the scDbs after single IP injection (Figure 1C). No evidence of toxicity or weight loss was observed during scDb administration (Figure 1D), suggesting that the scDbs are tolerable at this dosage in HIV-1-infected humanized mice on ART. After five weeks of ART, cohort I was euthanized to allow for viral reservoir measurement in tissues while ART was stopped for the mice of cohort II to allow for assessment of viral rebound.

**Figure 1.**
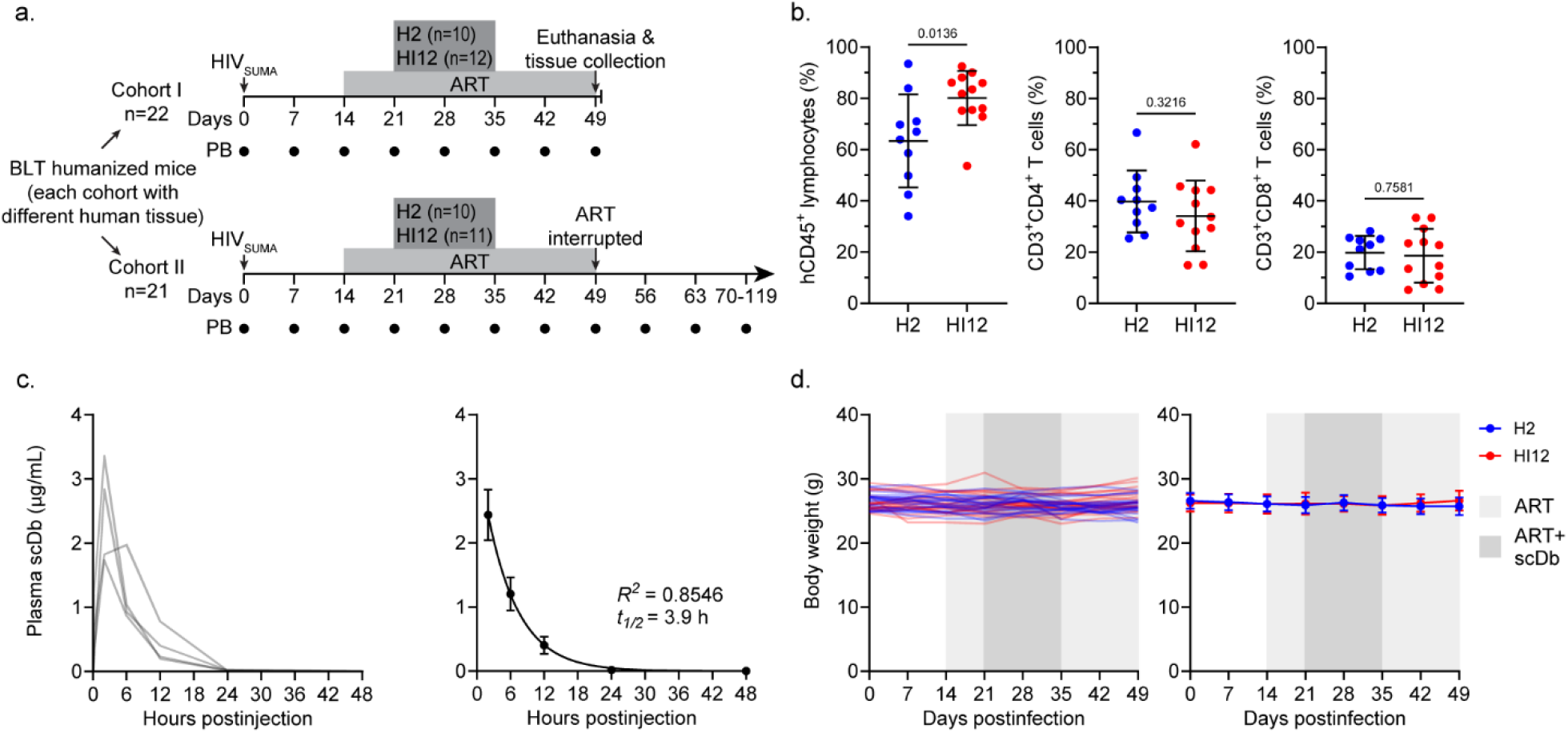
Study design and scDb tolerability. **a**, Overview of the study design. Two independent cohorts of humanized mice (n=22 for cohort I, n=21 for cohort II) were infected with HIV_SUMA_. Peripheral blood (PB) was collected weekly post-infection (black circles) for plasma and PBMCs. The mice of cohort II were euthanized on day 49 post-infection for tissue collection. ART was administered from day 14 to day 49 post-infection and H2 scDb or HI12 scDb were administered from day 21 to day 35 post-infection. **b**, Frequencies of hCD45^+^ lymphocytes, CD3^+^CD4^+^ T cells, and CD3^+^CD8^+^ T cells (mean, SD) in the peripheral blood 12 weeks after implantation of human fetal liver and thymus tissues. Significance determined by unpaired T-tests. **c**, scDb plasma pharmacokinetics in humanized mice. The mice received a 200 μg IP injection of H2 scDb and scDb plasma concentrations were assessed over time. Left: plasma scDb concentrations of the individual mice (n=4). Right: Average plasma scDb concentrations, mean, SE. Data were fitted to a single-phase exponential decay model (represented as line) and the predicted plasma half-life is indicated. **d**, Body weight of the humanized mice of cohort I reported in grams. Data shown for both individual mice (left) and as average values (right, mean, SD) for the H2 scDb (n=10, blue) and HI12 scDb (n=12, red) treatment groups. No significant differences between treatment groups at any time point nor significant changes in body weights between time points within either group as determined by two-way ANOVA followed by Šídák’s test for multiple comparisons.

To assess the ability of HI12 scDb to induce activation of peripheral blood T cells as indirect evidence of target recognition, we tracked the co-expression of two activation markers, CD38 and HLA-DR (Figure 2A,B,C). Following infection and viral replication, the frequency of activated (CD38^+^HLA-DR^+^) CD4^+^ and CD8^+^ T cells increased rapidly (Figure 2B,C), consistent with HIV-1-induced inflammation and the emergence of HIV-1-specific T cell responses. As expected, ART initiation resulted in a rapid decline of CD8^+^ T cell activation to near baseline levels and a gradual decline of CD4^+^ T cell activation (Figure 2B,C). Importantly, the introduction of HI12 scDb treatment after a week of ART significantly increased the frequency of CD38^+^HLA-DR^+^ CD8^+^ T cells relative to H2 scDb treatment by the end of dosing (Figure 2B). Although nonsignificant, HI12 scDb also resulted in a slightly elevated frequency of CD38^+^HLA-DR^+^ CD4^+^ T cells relative to H2 scDb (Figure 2C). Together, these results show HI12 scDb elevated activation of peripheral blood T cells in HLA-matched HIV-1-infected mice.

**Figure 2.**
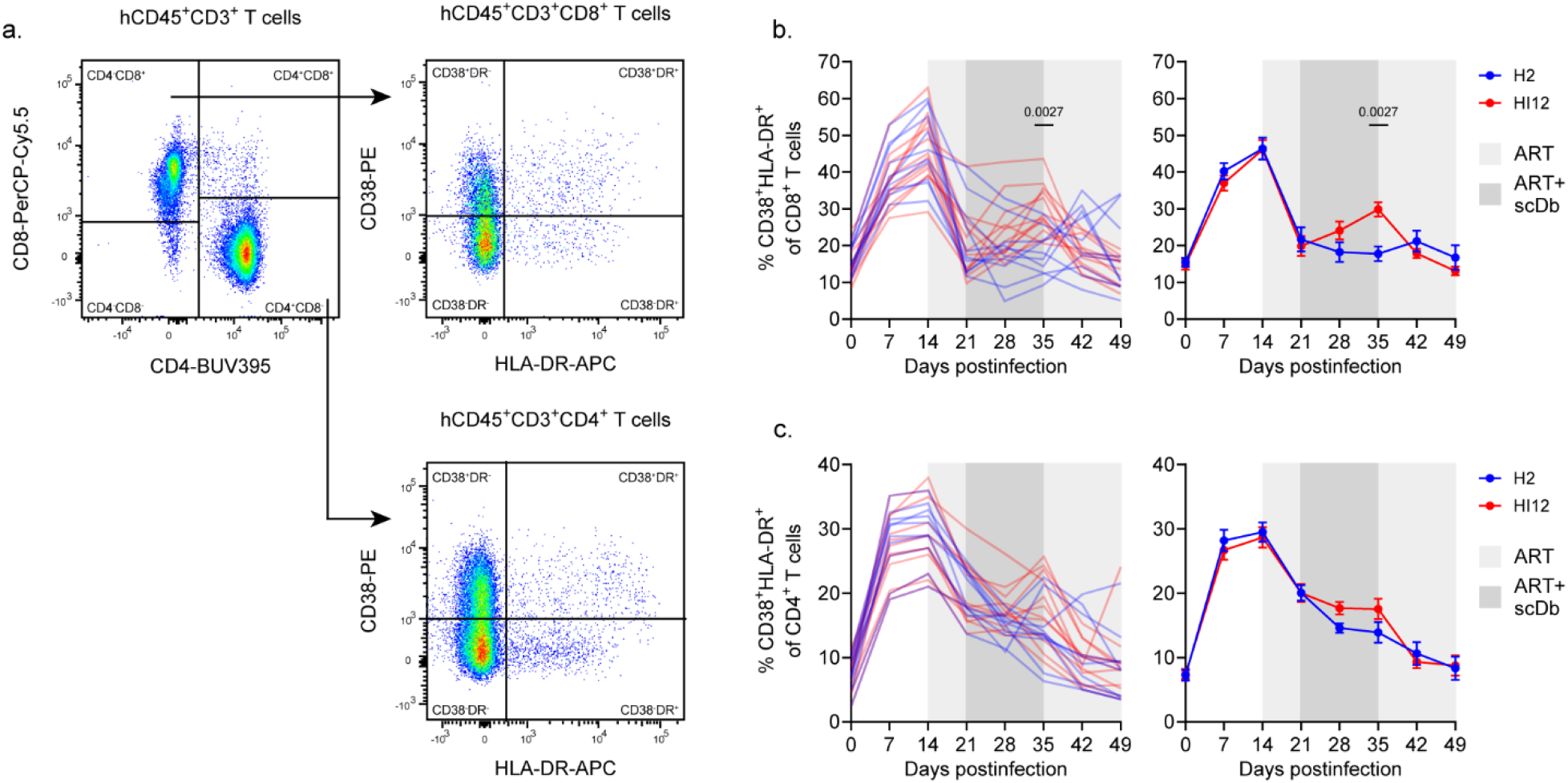
Immunological outcomes of scDb administration. **a**, Representative gating strategy for markers of T cell activation. Briefly, CD3^+^mCD45^-^hCD45^+^ T cells were segregated for analysis by CD4 and CD8 expression, and CD38^+^HLA-DR^+^ cells were quantified in within each T cell subset. **b, c**, Activation of peripheral blood T cells for cohort II. Data shown as the percent of CD38^+^HLA-DR^+^ cells within the CD3^+^CD4^-^CD8^+^ (**b**) or CD3^+^CD4^+^CD8^-^ (**c**) gates. Data shown for both individual mice (left) and as average values (right, mean, SE) for the H2 scDb (n=10, blue) and HI12 scDb (n=12, red) treatment groups. Significance determined by two-way ANOVA followed by Šídák’s test for multiple comparisons.

### HIV-1-specific TCR-mimic scDb significantly accelerated suppression after ART start and delayed time to viral rebound post ATI

Having established changes in CD8^+^ T cell activation following scDb administration, we evaluated the effect on HIV-1 plasma viremia during ART as well as upon ART interruption after viral suppression. Plasma viral load reached an average of 7.24x10^4^ HIV-1 RNA copies/mL by the time of ART initiation (two weeks postinfection) and declined to an average of 1.37x10^4^ HIV-1 RNA copies/mL by the time of scDb treatment initiation after one week of ART (Figure 3A,B). Neither at time of ART initiation nor at time of scDb treatment initiation did plasma viral load significantly differ between treatment and control groups (Figure 3B). Upon initiation of scDb treatment, plasma viral load declined more rapidly in the HI12 scDb-treated mice, reaching undetectable levels of plasma HIV-1 RNA 4.8 days earlier on average than H2 scDb-treated control mice (Figure 3A). By 4 weeks post-ART initiation, all mice had undetectable viral loads. Among the mice for which ART was interrupted to monitor viral rebound (cohort II), rebound viremia became detectable ranging from one to four weeks after cessation of ART (Figure 3C). Importantly, the mice treated with HI12 scDb exhibited an average delay in viral rebound of 11.0 days when compared to controls treated with H2 scDb (Figure 3C,D). Notably, we were unable to detect HIV-1 RNA in the plasma of one mouse in the HI12 scDb-treated group for up to ten weeks after cessation of ART. To determine whether rebound viremia represented escape from the epitope targeted by the HI12 scDb (ILKEPVHGV, Pol 464-472), we recovered *pol* sequences from plasma at the earliest time points with detectable viral load following ART cessation. However, we did not find considerable evidence of selection for mutations in the target epitope in the predominant viral clones contributing to rebound (data not shown). The faster time to viral suppression and the delayed time to viral rebound upon ART interruption observed in the HI12 scDb-treated mice are consistent with enhanced CD8^+^ T cell mediated clearance of HIV-1-infected CD4^+^ T cells during scDb administration.

**Figure 3.**
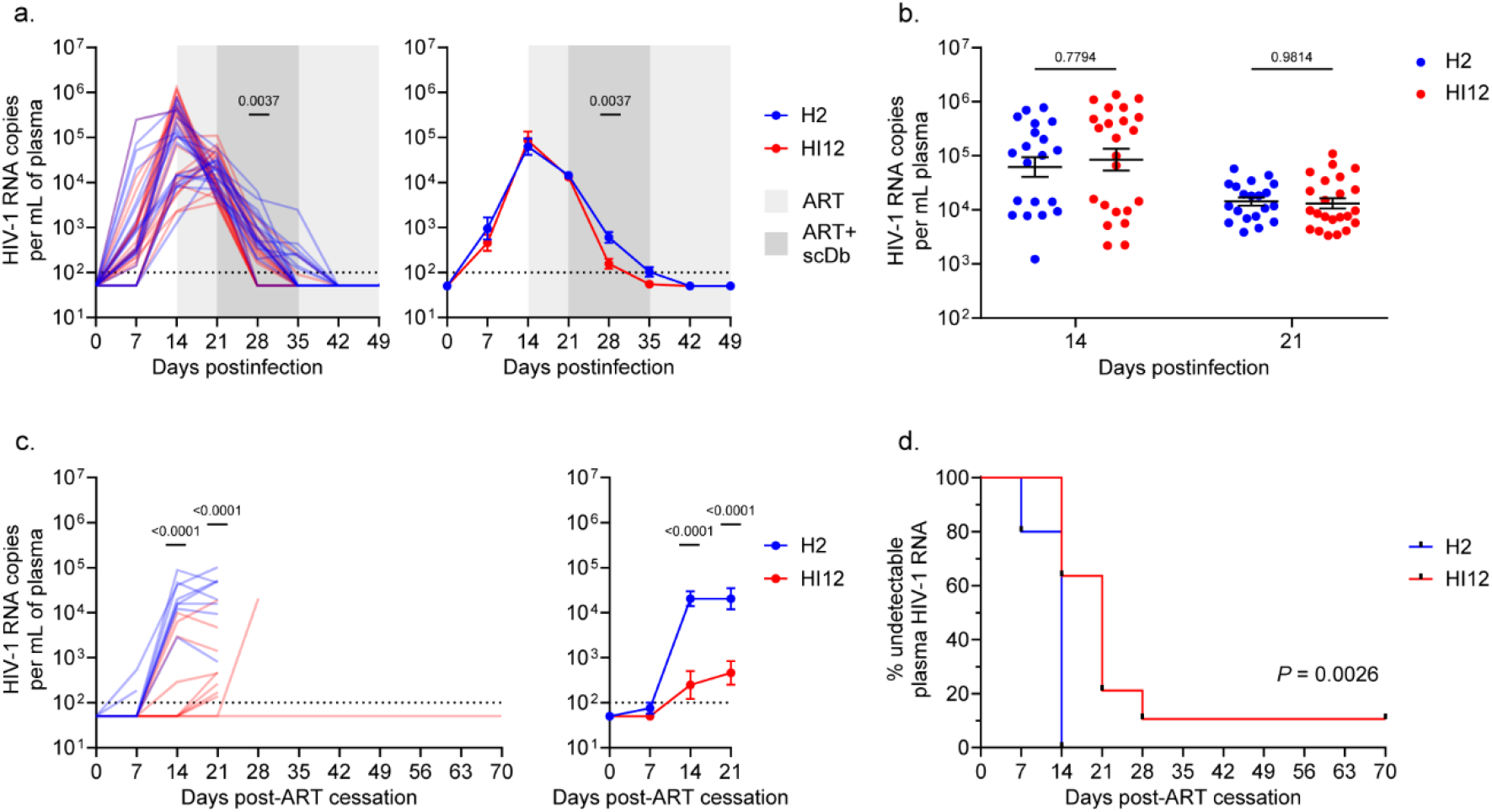
scDb-induced changes in HIV-1 plasma viremia. **a**, HIV-1 plasma RNA before and during ART administration (day 0 to day 49 post-infection) for cohorts I and II. HIV-1 RNA copies/mL plasma shown for both individual mice (left) and as average values (right, mean, SE) for the H2 scDb (n=10 for cohort I, n=10 for cohort II, blue) and HI12 scDb (n=12 for cohort I, n=11 for cohort II, red) treatment groups. Significance determined by two-way ANOVA followed by Šídák’s test for multiple comparisons. **b**, Comparison of HIV-1 plasma RNA between treatment groups at initiation of ART (day 14) and at initiation of scDb administration (day 21) as in (**a**). **c**, HIV-1 plasma RNA rebound following ART cessation (day 49 to day 119 post-infection) for cohort II. HIV-1 RNA copies/mL plasma shown for both individual mice (left) and as average values (right, mean, SE) for the H2 scDb (n=10, blue) and HI12 scDb (n=11, red) treatment groups. Significance determined by two-way ANOVA followed by Šídák’s test for multiple comparisons. **d**, Survival analysis of mice with undetectable HIV-1 plasma RNA (<100 copies/mL) post-ART cessation (day 49 to day 119 post-infection). Significance determined by log-rank test.

### HIV-1-specific TCR-mimic scDb reduced HIV-1 DNA burden in lymphoid tissues

To validate that the observed delay in viral rebound is linked to a reduction in HIV-1-infected cells, we evaluated HIV-1 DNA burden in splenic and lymph node (LN) tissues at five weeks after ART initiation when full viral suppression was observed. We utilized two complimentary methods for quantifying HIV-1 proviruses: droplet digital PCR and HIV-1 DNAscope in situ hybridization. First, we analyzed splenocytes using the intact proviral DNA assay (IPDA)^16^, a highly sensitive droplet digital PCR assay for quantifying HIV-1 proviruses. The IPDA measures both sequence-intact HIV-1 proviruses, which are expected to be the source of replication competent virus responsible for viral rebound, in addition to proviruses with common defects, including large internal deletions or APOBEC3G-mediated hypermutation — which constitute the vast majority of proviruses in the latent reservoir^16^. The IPDA was performed using DNA extracted from splenocytes of the cohort I mice (n=10 for H2 scDb, n=11 for HI12 scDb). We found a 2.9-fold reduction in total HIV-1 proviruses in the HI12 scDb treatment group when compared to controls treated with H2 scDb (Figure 4A). Importantly and consistent with the delay of viral rebound, IPDA results showed a 4.1-fold reduction in intact proviruses. To further corroborate our results by an independent assay on a different lymphoid tissue, we used anti-CD4 staining together with DNAscope in situ hybridyzation^17^, enabling visualization and quantification of HIV-1 DNA at single-copy and single-cell resolution. This technique enabled measurement of HIV-1 proviruses in paraformaldehyde (PFA)-fixed lymph node tissue sections from a subset of cohort I mice (n=6 for H2 scDb, and n=5 for HI12 scDb). CD4^+^ cells harboring HIV-1 proviruses were detected in all LN tissues analyzed (Figure 4B,C). Results showed a 5.7-fold reduction in HIV-1 total proviral DNA per lymph node CD4^+^ cell in HI12 scDb-treated mice as compared to control H2 scDb-treated mice (Figure 4C). Taken together, HI12 scDb treatment reduced HIV-1 DNA burden in lymphoid tissues after ART suppression. This remarkable decrease likely underlies the observed delay in viral rebound as a consequence of enhanced CD8^+^ T cell-mediated clearance of infected cells during early ART.

**Figure 4.**
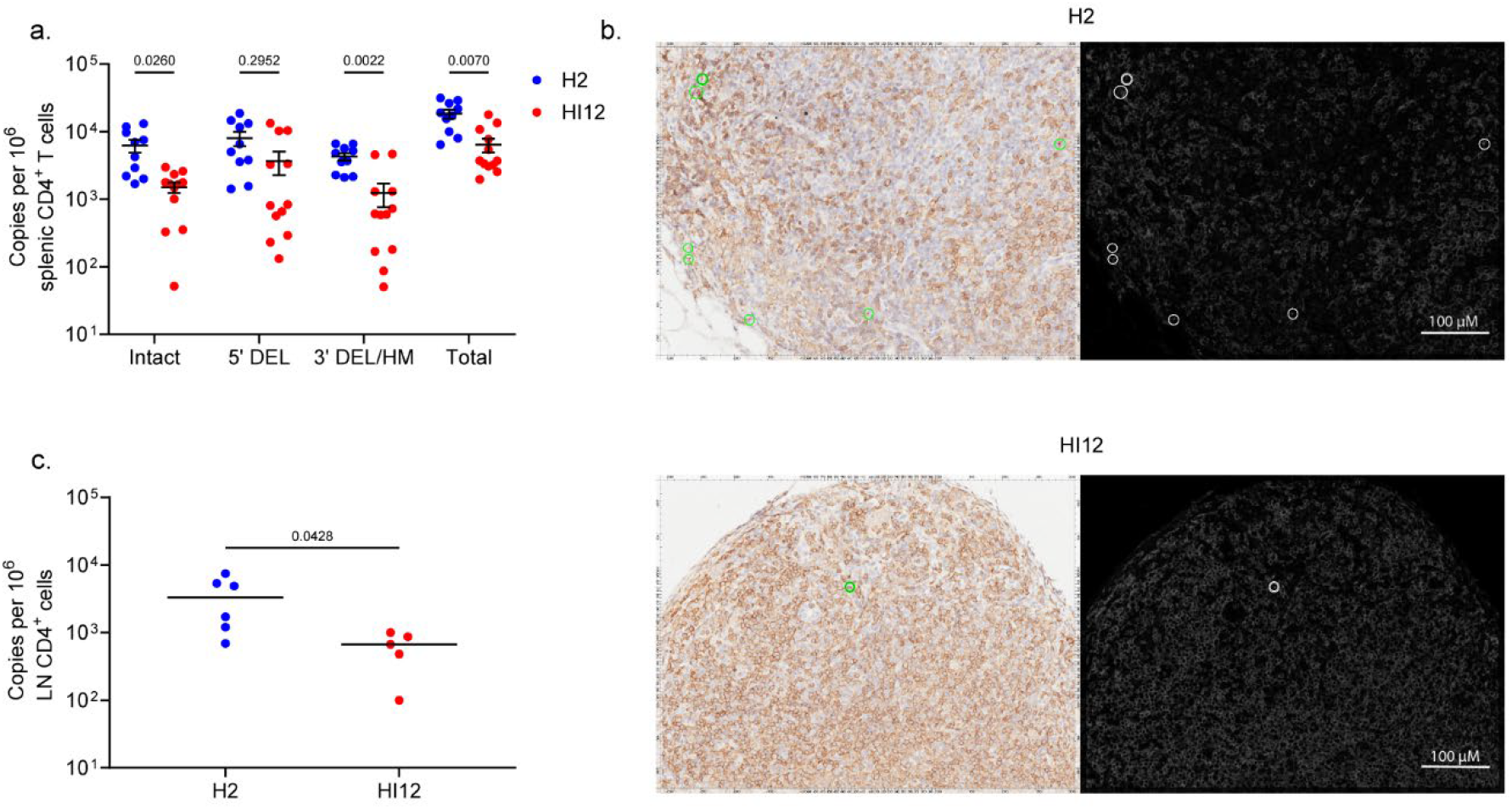
scDb-induced changes in HIV-1 cellular reservoirs. **a**, Frequencies of splenic CD4^+^ T cells harboring intact, 5’ deleted, 3’ deleted/hypermutated, or total HIV-1 provirus (cohort I) quantified by IPDA. Data reported as HIV-1 copies per million splenic CD4^+^ T cells (mean, SE, n=10 for H2 scDb, n=12 for HI12 scDb). Significance determined by two-way ANOVA followed by Šídák’s test for multiple comparisons. **b**, Representative images of CD4^+^ T cells (brown) harboring HIV-1 DNA (red) in PFA-fixed LN tissues from an H2 scDb-treated mouse (top) and an HI12-treated mouse (bottom) of cohort I. Scale bar =100μm. **c**, Frequencies of LN CD4^+^ T cells harboring HIV-1 provirus (cohort I) quantified by DNAscope. Data reported as HIV-1 copies per million LN CD4^+^ T cells (median, n=6 for H2 scDb, n=5 for HI12 scDb). Significance determined by unpaired T-test.

## Discussion

Here we provide the first in vivo evidence that a peptide-HLA-targeted bispecific antibody can measurably shrink the intact HIV reservoir and delay rebound after antiretroviral therapy interruption. By redirecting CD8^+^ T cells—irrespective of their native specificity—toward a single conserved Pol-derived epitope displayed on infected cells, the HI12 TCR-mimic diabody bypasses the need for Env expression or for vaccine-primed HIV-specific T-cell responses. In the setting of acute infection, initial HIV-1-specific CD8^+^ T cell responses play a substantial role in shaping long-term reservoir dynamics and disease progression outcomes^18,19^. Our data is consistent with CD8^+^ T cell-mediated antiviral pressure, as we found that early treatment with HI12 scDb could alter viral decay kinetics (Figure 3A,B) with a subsequent reduction of HIV-1 DNA burden (Figure 4A-C) and a corresponding delay in viral rebound after ART interruption (Figure 3C,D).

TCR-based immune pressure is known to select for HIV-1 isolates with mutations in the corresponding epitope during active replication, resulting in viral escape and diminished efficacy of subsequent responses directed against that epitope^20–22^. In the context of the present study, it is important to highlight that scDb administration occurred during ART where the presence of antivirals would prevent HIV-1 replication and thus the effects of any selective pressure exerted by CD8^+^ T cells may be less apparent. Targeting of multiple distinct conserved HIV-1 epitopes by CD8^+^ T cell pressure may prove more efficacious at limiting or delaying the emergence of viral escape mutations than single epitope approaches^22–24^. As such, future studies could explore coadministration of HI12 scDb with TCR-mimic scDbs directed against alternative HIV epitopes both during early ART and at viral rebound after ART interruption.

While our strategy was based on peptide:HLA-directed constructs, a complementary approach for the design of HIV-1-targeting bispecific antibodies could have included antibody modalities recognizing HIV-1 Env^8,9,13,25,26^. On the one hand, Env-directed antibodies would not be restricted to clinical use in PWH with the corresponding HLA allele. However, given the high degree of sequence diversity in HIV-1 Env and the narrow breadth of Env binding attainable by most antibody clones, peptide:HLA-directed approaches as shown here may have wider application by targeting intracellular viral proteins such as Gag or Pol with a high degree of conservation across viral isolates and in which escape mutations are more likely to result in viral fitness costs^27,28^. In line with this, the HI12 epitope is highly conserved across clade B HIV-1 (96.0% of isolates with an identical amino acid match or a single amino acid substitution, Los Alamos HIV sequence database, n=2396). Our prior study demonstrated that single amino substitutions in the target epitope of TCR-mimic scDbs are often tolerable for scDb binding^12^, which may enable recognition of a wider range of viral isolates. Furthermore, HIV-1 Env on infected cell surfaces is greatly diminished or absent during suppressive ART, while peptide:MHC complexes on infected cell surfaces are more stable and can persist longer than transient viral antigen expression^29^. We have also previously identified that the sequence landscape of the latent reservoir includes a significant number of defects in the *env* gene^30,31^. To this end, peptide:HLA-directed bispecific antibodies may also have greater utility in eliminating cells containing proviruses with major splice donor defects or large 3’ deletions which may limit Env expression but still allow for production of Gag and Pol proteins^32–34^. Consistent with this idea, HI12 scDb, which recognizes a Pol-derived epitope, resulted in more significant reductions in cells containing intact proviruses and proviruses with 3’ deletions or hypermutation than those with 5’ deletions (Figure 4A). Of the three categories of proviruses differentiated by the IPDA, those containing 5’ deletions are most likely to have defects affecting Pol expression and are therefore more likely to restrict Pol-dependent targeting modalities.

The HI12 scDb described in this study relies on an anti-CD3 scFv to recruit CD8^+^ T cells to infected targets and activate cytolytic and cytokine secretion pathways. This modality could be substituted with scFvs recognizing activating receptors on other immune effector cell types to elicit broader anti-HIV-1 responses. For example, scDbs with anti-CD16 modalities have been shown to elicit HIV-1-specific NK cell responses in both cell culture and humanized mouse model systems^10,11,13^. Exploration of HIV-1 therapeutics that engage innate immune effectors such as NK cells in humanized mice will require specialized models capable of supporting the development of these cell lineages^13,35–37^.

A clear limitation of this strategy was highlighted by the limited pharmacokinetics of the scDb construct observed after infusion (Figure 1C). The short half-life of the TCR-mimic scDbs required necessitate daily injections or continuous infusion to maintain effective therapeutic concentrations, potentially complicating clinical application. Future iterations of TCR-mimic scDbs will need to be enhanced through protein modifications that prolong their in vivo half-life^38,39^. It also remains to be determined if this strategy could be tailored to different HLAs and/or target multiple epitopes concurrently. An additional safety consideration for clinical application of TCR-mimic scDbs is the potential for nonspecific T cell activation and cytokine release through CD3 engagement. Neither our previous in vitro characterization of TCR-mimic scDbs^12^ nor the present study document compelling evidence of broad off-target T cell activation. Finally, while our infection model using a single T/F HIV-1 clone (HIV_SUMA_) containing the HI12 epitope allowed us to demonstrate a proof of concept that HLA-directed strategies can enhance HIV-1 clearance during early ART, future studies will need to test the activity of TCR-mimic scDbs against more diverse viral populations, as is present in chronic infection.

Lymphoid tissues are the primary locations of HIV-1 replication during untreated infection and contain the majority of infected CD4^+^ T cells in the body after ART^40^. Additionally, recent studies have found evidence of reduced antiretroviral drug concentrations and low level HIV-1 RNA expression in lymphoid tissue sites^41–45^, suggesting that suppression of viral replication at these sites may be incomplete. Therefore, infected cells residing in lymphoid tissues are expected to contribute to viral rebound upon treatment interruption and therapeutic approaches to reservoir reduction must demonstrate efficacy at these anatomical sites. However, several factors make elimination of HIV-1-infected cells more challenging in tissue sites than in peripheral blood, including potentially reduced bioavailability of the therapeutic, limited infiltration of cytotoxic effectors, and immunomodulatory effects of the tissue microenvironment. Importantly, we show here that HI12 scDb is capable of reducing HIV-1 DNA burden in secondary lymphoid tissues (Figure 4A-C) in addition to resulting in a delay of viral rebound.

Our data also opens the potential to combine TCR-mimic scDbs with additional antiviral strategies such as latency reversing agents to drive expression and presentation of viral antigens during ART. Although conventional latency reversal strategies often fail to induce robust and sustained viral expression^46^, more recently developed agents such as activators of the noncanonical NF-κB signaling pathway and the IL-15 superagonist, N-803, have demonstrated improved efficacy in preclinical animal models of HIV infection^47–49^ and therefore may enable greater TCR mimic scDb-mediated clearance of infected cells. Overall, our study not only validates HLA-restricted redirection of CD8^+^ T cell responses as an effective stand-alone anti-HIV strategy but also establishes a benchmark for future regimens with multi-epitope TCR-mimic scDbs together with complementary immune effectors in HIV-cure directed interventions.

## Materials and Methods

### Generation of HLA specific humanized mice

BLT mice were generated as previously described^13,50–52^, in accordance with the Wistar Institute Animal Care and Research Committee regulations (protocol #201360). Briefly, 6-to 8-week-old female NSG (NOD.Cg-Prkdc^scid^ Il2rg^tm1Wjl^/SzJ, Jackson Laboratory) mice were pretreated with busulfan at 30mg/kg and were then implanted with human fetal thymic tissue fragments and fetal liver tissue fragments under the murine renal capsule. Following the surgery, mice were injected via the tail vein with CD34^+^ hematopoietic stem cells isolated from human fetal liver tissues. Human fetal liver and thymus tissues were procured from Advanced Bioscience Resources and prescreened for the HLA-A^*^02:01 allele by sequence specific PCR. Briefly, genomic DNA from human tissues was extracted by Monarch Genomic DNA Purification Kit (NEB, # T3010S). The HLA-A locus sequence was determined with the Olerup SSP HLA-A genotype kit (CareDx Inc. #101.401-48). Each of the two independent cohorts of BLT mice generated were reconstituted with tissues from a different HLA-A^*^02:01^+^ donor. 12 weeks post-surgery, human immune cell reconstitution in peripheral blood was determined using a FACSymphony flow cytometer (BD Biosciences) using the following antibodies: mCD45-AF700, hCD45-FITC, hCD3-BUV805, hCD4-BUV395, hCD8-PerCP-Cy5.5 and Fixable Viability Stain 510 (catalog# 560510, 555482, 612895, 563550, 565310, and 564406, respectively; BD Biosciences). Data was analyzed with FlowJo.

### HIV-1 infection, ART suppression, and scDb administration

Each cohort of humanized mice was randomly divided into two subgroups to receive treatment with H2 or HI12 scDb (cohort I, n=10 for H2 and n=12 for HI12; cohort II, n=10 for H2 and n=11 for HI12). The mice were infected intravenously with 1×10^4^ TCID_50_ (50% tissue culture infectious dose) of T/F virus HIV_SUMA_. Peripheral blood was collected weekly for assessment of peripheral blood T cell activation and plasma viral load. Then, 2 weeks post infection, the mice were placed on a diet combined with ART (1,500 mg/kg of emtricitabine, 1,560 mg/kg of tenofovir disoproxil fumarate, and 600 mg/kg of raltegravir). Starting at 3 weeks postinfection and ending at 5 weeks postinfection, the mice were given daily intraperitoneal injections of 200 μg (approximately 7 mg/kg) of either H2 or HI12 scDb. At 7 weeks postinfection, the mice of cohort I were euthanized and tissues were collected for DNAscope and IPDA, and the mice of cohort II were returned to a diet without ART. Plasma viral loads of the cohort II mice were monitored weekly following ART cessation and the mice were euthanized following detection of viral rebound.

### scDb plasma pharmacokinetic measurement

scDb plasma decay in the humanized mice was measured as previously described^13^. Single 200 μg IP injections of H2 scDb were administered to a group of BLT mice (n=4, independent donor tissue from those used for cohorts I and II). scDb plasma concentrations were assessed at 0-, 0.2-, 2-, 6-, 12-, 24-, and 48-hours post-injection by ELISA. Briefly, biotinylated peptide:HLA-A^*^02:01 monomer corresponding to the H2 epitope (Fred Hutchinson Immune Monitoring Lab, 1 μg/mL) in BAE blocking buffer (PBS, 0.5% bovine serum albumin, 0.1% sodium azide) were added to an EvenCoat streptavidin-coated plate (R&D Systems) and incubated at 4 °C for 16 h. The plate was washed with 1× TBS-T (1× tris-buffered saline (TBS) + 0.05% Tween-20) using a BioTek 405 plate washer and plasma samples diluted 1:2 in BAE were plated. A standard curve of H2 scDb prepared in a 1:2 mixture of pre-injection plasma:BAE was plated in parallel. After 16h of incubation at 4 °C, the plate was washed as before and HRP (horse radish peroxidase) Protein L secondary (Thermo Fisher Scientific, catalog no. 32420, 0.5 μg/ml in BAE) was plated and incubated at room temperature for 1 h. The plate was washed and developed in TMB (3,3′,5,5′- tetramethylbenzidine) for 5 min at room temperature before stopping in 1 N sulfuric acid and reading the optical density at 450 nm (OD450). A dose-response model was fitted to the standard curve measurements and used to calculate H2 scDb plasma concentrations. The 2-, 6-, 12-, 24-, and 48-hours post-injection H2 scDb plasma concentrations were then fitted to a single-phase exponential decay model to determine the predicted plasma half-life.

### T cell activation measurement

Peripheral blood mononuclear cells (PBMCs) were stained using the following antibodies (1:50 dilution): mCD45-AF700, hCD45-FITC, hCD3-BUV805, hCD4-BUV395, hCD8-PerCP-Cy5.5, HLA-DR-APC, hCD38-PE, and Fixable Viability Stain 510 (catalog nos. 560510, 555482, 612895, 563550, 565310, 340691, 555460, and 564406, respectively; BD Biosciences). Data were collected on a FACSymphony flow cytometer (BD Biosciences). Data were analyzed with FlowJo.

### Plasma viral load measurement

Plasma viral loads were measured as previously described^13,50–54^. Briefly, viral RNA was extracted using a QIAamp Viral RNA Mini kit (QIAGEN). The pVLs were determined using reverse transcription quantitative PCR (RT-qPCR) on a C1000 Thermal Cycler and the CFX96 Real-Time system (BioRad) with the TaqMan Fast Virus 1-Step Master Mix (Life Technologies).

### DNAscope and immunohistochemistry

To detect HIV-1 DNA and T cell marker CD4 simultaneously, a sequential combination of DNAscope in situ hybridization with immunohistochemical staining (IHCS) was conducted. DNAscope was conducted according to modified Advanced Cell Diagnostics (ACD)’s protocol as previously described^55^. Briefly, after deparaffinization and rehydration, tissue sections were subjected to antigen retrieval through boiling for 15 min in RNAscope Target Retrieval Reagent solution (Cat. no. 322000, ACD, Farmington, UT, USA) and treating with RNAscope Protease Plus (cat. no. 322330, ACD) at 40°C for 15 min. For HIV DNA detection, tissue sections were hybridized with V-HIV1-CladeB-sense probe (cat. no. 425531, ACD) at 40°C for 2 hours, respectively. The detection and amplification of signals were carried out using RNAscope 2.5 HD assay-RED kit (Cat. no. 322360, ACD). After the completion of DNAscope, the tissue sections were immediately subjected to IHCS. After blocked with a blocking buffer consisting of 5% bovine serum albumin, tissue sections were incubated with rabbit anti-human CD4 antibody (1:200 dilution; EPR6855; Abcam) at 4°C overnight. DAB (3,3’-Diaminobenzidine) was used to visualize antibody staining signals using Envision GI2 Doublestain System Rabbit/Mouse kit (Dako, K5361) based on the manufacturer’s manual. After counterstained with hematoxylin, tissue sections were digitized with Aperio CS2 Scanscope and were analyzed using Aperio’s Spectrum Plus analysis program (version 9.1; Aperio ePathology Solutions). The frequency of vDNA positive cells in CD4^+^ T cells were manually counted.

### Intact proviral DNA assay

The IPDA was used to quantify HIV-1 proviral DNA in humanized mouse-derived splenocytes as previously described^13^. Briefly, single-cell suspensions of splenocytes were generated using the gentleMACS Octo Dissociator (Miltenyi Biotec) and genomic DNA was extracted from the resulting splenocyte suspensions. Each IPDA ddPCR reaction was performed in parallel with a copy reference/shearing correction (RPP30 gene) ddPCR reaction which quantifies input human cell equivalents and DNA shearing^16^. Input human CD4^+^ T cell equivalents were calculated based on flow cytometry staining of the splenocyte samples as follows: (Input human RPP30 cell equivalents) × (percentage hCD3^+^CD4^+^ cells of total hCD45^+^ cells). Shearing corrections were applied to the IPDA data and the final results are reported as copies per 10^6^ human spleen-derived CD4^+^ T cells.

### Statistical analysis

Statistical analyses used in the present study include Student’s t-tests (Figures 1B, 4C), nonlinear regression models (Figure 1C), analysis of variance (ANOVA) with multiple comparisons (Figures 1D, 2B-C, 3A-C, 4A), and Kaplan–Meier survival analysis (Figure 3D). All statistical analyses were performed using GraphPad Prism. Data distribution was assumed to be normal for all analyses but this was not formally tested. Two-sided P < 0.05 was considered significant for all tests.

## Data availability

The data that supports the findings of this study are available from the corresponding authors.

## Acknowledgments

This work was supported by the BEAT-HIV (UM1 AI164570), I4C (UM1AI164556), iCURE (UM1AI191272), and DARE (UM1AI164560) NIH Martin Delaney Collaboratories, and by the Penn Center for AIDS Research (P30AI045008). L.J.M. is supported by the Robert I. Jacobs Fund of the Philadelphia Foundation and the Herbert Kean, M.D., Family Professorship. R.F.S. is supported by the Howard Hughes Medical Institute.

## References

1. Perelson, A. S. et al. Decay characteristics of HIV-1-infected compartments during combination therapy. Nature 387, 188–91 (1997).

2. Vittinghoff, E. et al. Combination antiretroviral therapy and recent declines in AIDS incidence and mortality. J. Infect. Dis. 179, 717–20 (1999).

3. Chun, T. W. et al. In vivo fate of HIV-1-infected T cells: quantitative analysis of the transition to stable latency. Nat. Med. 1, 1284–90 (1995).

4. Chun, T. W. et al. Quantification of latent tissue reservoirs and total body viral load in HIV-1 infection. Nature 387, 183–8 (1997).

5. Siliciano, J. D. et al. Long-term follow-up studies confirm the stability of the latent reservoir for HIV-1 in resting CD4+ T cells. Nat. Med. 9, 727–8 (2003).

6. Peluso, M. J. et al. Differential decay of intact and defective proviral DNA in HIV-1-infected individuals on suppressive antiretroviral therapy. JCI insight 5, e132997 (2020).

7. McMyn, N. F. et al. The latent reservoir of inducible, infectious HIV-1 does not decrease despite decades of antiretroviral therapy. J. Clin. Invest. 133, e171554 (2023).

8. Sung, J. A. M. et al. Dual-Affinity Re-Targeting proteins direct T cell-mediated cytolysis of latently HIV-infected cells. J. Clin. Invest. 125, 4077–90 (2015).

9. Pegu, A. et al. Activation and lysis of human CD4 cells latently infected with HIV-1. Nat. Commun. 6, 8447 (2015).

10. Pollara, J. et al. Redirection of Cord Blood T Cells and Natural Killer Cells for Elimination of Autologous HIV-1-Infected Target Cells Using Bispecific DART® Molecules. Front. Immunol. 11, 713 (2020).

11. Ramadoss, N. S. et al. Enhancing natural killer cell function with gp41-targeting bispecific antibodies to combat HIV infection. AIDS 34, 1313–1323 (2020).

12. Sengupta, S. et al. TCR-mimic bispecific antibodies to target the HIV-1 reservoir. Proc. Natl. Acad. Sci. U. S. A. 119, e2123406119 (2022).

13. Board, N. L. et al. Bispecific antibodies promote natural killer cell-mediated elimination of HIV-1 reservoir cells. Nat. Immunol. 25, 462–470 (2024).

14. Hsiue, E. H.-C. et al. Targeting a neoantigen derived from a common TP53 mutation. Science 371, eabc8697 (2021).

15. Douglass, J. et al. Bispecific antibodies targeting mutant RAS neoantigens. Sci. Immunol. 6, eabd5515 (2021).

16. Bruner, K. M. et al. A quantitative approach for measuring the reservoir of latent HIV-1 proviruses. Nature 566, 120–125 (2019).

17. Deleage, C. et al. Defining HIV and SIV Reservoirs in Lymphoid Tissues. Pathog. Immun. 1, 68 (2016).

18. Goonetilleke, N. et al. The first T cell response to transmitted/founder virus contributes to the control of acute viremia in HIV-1 infection. J. Exp. Med. 206, 1253–72 (2009).

19. Streeck, H. et al. Human immunodeficiency virus type 1-specific CD8^+^ T-cell responses during primary infection are major determinants of the viral set point and loss of CD4+ T cells. J. Virol. 83, 7641–8 (2009).

20. Leslie, A. J. et al. HIV evolution: CTL escape mutation and reversion after transmission. Nat. Med. 10, 282–9 (2004).

21. Goulder, P. J. R. & Watkins, D. I. HIV and SIV CTL escape: implications for vaccine design. Nat. Rev. Immunol. 4, 630–640 (2004).

22. Deng, K. et al. Broad CTL response is required to clear latent HIV-1 due to dominance of escape mutations. Nature 517, 381–5 (2015).

23. Garcia, V. & Regoes, R. R. The Effect of Interference on the CD8(+) T Cell Escape Rates in HIV. Front. Immunol. 5, 661 (2014).

24. Garcia, V., Feldman, M. W. & Regoes, R. R. Investigating the Consequences of Interference between Multiple CD8^+^ T Cell Escape Mutations in Early HIV Infection. PLoS Comput. Biol. 12, e1004721 (2016).

25. Sloan, D. D. et al. Targeting HIV Reservoir in Infected CD4 T Cells by Dual-Affinity Retargeting Molecules (DARTs) that Bind HIV Envelope and Recruit Cytotoxic T Cells. PLoS Pathog. 11, e1005233 (2015).

26. Brozy, J. et al. Antiviral Activity of HIV gp120-Targeting Bispecific T Cell Engager Antibody Constructs. J. Virol. 92, e00491–18 (2018).

27. Rihn, S. J. et al. Extreme genetic fragility of the HIV-1 capsid. PLoS Pathog. 9, e1003461 (2013).

28. Gaiha, G. D. et al. Structural topology defines protective CD8^+^ T cell epitopes in the HIV proteome. Science 364, 480–484 (2019).

29. Lazaro, E. et al. Variable HIV peptide stability in human cytosol is critical to epitope presentation and immune escape. J. Clin. Invest. 121, 2480–92 (2011).

30. Ho, Y.-C. et al. Replication-competent noninduced proviruses in the latent reservoir increase barrier to HIV-1 cure. Cell 155, 540–51 (2013).

31. Bruner, K. M. et al. Defective proviruses rapidly accumulate during acute HIV-1 infection. Nat. Med. 22, 1043–9 (2016).

32. Pollack, R. A. et al. Defective HIV-1 Proviruses Are Expressed and Can Be Recognized by Cytotoxic T Lymphocytes, which Shape the Proviral Landscape. Cell Host Microbe 21, 494-506.e4 (2017).

33. Imamichi, H. et al. Defective HIV-1 proviruses produce viral proteins. Proc. Natl. Acad. Sci. U. S. A. 117, 3704–3710 (2020).

34. White, J. A. et al. Clonally expanded HIV-1 proviruses with 5’-leader defects can give rise to nonsuppressible residual viremia. J. Clin. Invest. 133, e165245 (2023).

35. Herndler-Brandstetter, D. et al. Humanized mouse model supports development, function, and tissue residency of human natural killer cells. Proc. Natl. Acad. Sci. U. S. A. 114, E9626–E9634 (2017).

36. Sungur, C. M. et al. Human NK cells confer protection against HIV-1 infection in humanized mice. J. Clin. Invest. 132, e162694 (2022).

37. Kim, J. T., Bresson-Tan, G. & Zack, J. A. Current Advances in Humanized Mouse Models for Studying NK Cells and HIV Infection. Microorganisms 11, 1984 (2023).

38. Stork, R., Campigna, E., Robert, B., Müller, D. & Kontermann, R. E. Biodistribution of a bispecific single-chain diabody and its half-life extended derivatives. J. Biol. Chem. 284, 25612–9 (2009).

39. Unverdorben, F., Färber-Schwarz, A., Richter, F., Hutt, M. & Kontermann, R. E. Half-life extension of a single-chain diabody by fusion to domain B of staphylococcal protein A. Protein Eng. Des. Sel. 25, 81–8 (2012).

40. Wong, J. K. & Yukl, S. A. Tissue reservoirs of HIV. Curr. Opin. HIV AIDS 11, 362–70 (2016).

41. Fletcher, C. V et al. Persistent HIV-1 replication is associated with lower antiretroviral drug concentrations in lymphatic tissues. Proc. Natl. Acad. Sci. U. S. A. 111, 2307–12 (2014).

42. Burgunder, E. et al. Antiretroviral Drug Concentrations in Lymph Nodes: A Cross-Species Comparison of the Effect of Drug Transporter Expression, Viral Infection, and Sex in Humanized Mice, Nonhuman Primates, and Humans. J. Pharmacol. Exp. Ther. 370, 360– 368 (2019).

43. Thompson, C. G. et al. Heterogeneous antiretroviral drug distribution and HIV/SHIV detection in the gut of three species. Sci. Transl. Med. 11, eaap8758 (2019).

44. Devanathan, A. S. et al. Antiretroviral Penetration and Drug Transporter Concentrations in the Spleens of Three Preclinical Animal Models and Humans. Antimicrob. Agents Chemother. 64, e01384–20 (2020).

45. Fletcher, C. V et al. Persistent HIV transcription and variable antiretroviral drug penetration in lymph nodes during plasma viral suppression. AIDS 36, 985–990 (2022).

46. Tsai, P. et al. In vivo analysis of the effect of panobinostat on cell-associated HIV RNA and DNA levels and latent HIV infection. Retrovirology 13, 36 (2016).

47. Bobardt, M. et al. The inhibitor apoptosis protein antagonist Debio 1143 Is an attractive HIV-1 latency reversal candidate. PLoS One 14, e0211746 (2019).

48. Nixon, C. C. et al. Systemic HIV and SIV latency reversal via non-canonical NF-κB signalling in vivo. Nature 578, 160–165 (2020).

49. McBrien, J. B. et al. Robust and persistent reactivation of SIV and HIV by N-803 and depletion of CD8^+^ cells. Nature 578, 154–159 (2020).

50. Yuan, Z. et al. Recapitulating Cross-Species Transmission of Simian Immunodeficiency Virus SIVcpz to Humans by Using Humanized BLT Mice. J. Virol. 90, 7728–39 (2016).

51. Yuan, Z., Kang, G., Daharsh, L., Fan, W. & Li, Q. SIVcpz closely related to the ancestral HIV-1 is less or non-pathogenic to humans in a hu-BLT mouse model. Emerg. Microbes Infect. 7, 59 (2018).

52. Yuan, Z. et al. Human galectin-9 promotes the expansion of HIV reservoirs in vivo in humanized mice. AIDS 37, 571–577 (2023).

53. Yuan, Z. et al. Controlling Multicycle Replication of Live-Attenuated HIV-1 Using an Unnatural Genetic Switch. ACS Synth. Biol. 6, 721–731 (2017).

54. Yuan, Z., Kang, G., Lu, W. & Li, Q. Reactivation of HIV-1 proviruses in immune-compromised mice engrafted with human VOA-negative CD4+ T cells. J. virus Erad. 3, 61–65 (2017).

55. Ko, A. et al. Macrophages but not Astrocytes Harbor HIV DNA in the Brains of HIV-1-Infected Aviremic Individuals on Suppressive Antiretroviral Therapy. J. Neuroimmune Pharmacol. 14, 110–119 (2019).

